# Relationship between personality and cognitive traits in domestic rabbits (*Oryctolagus cuniculus*)

**DOI:** 10.1101/2020.10.12.336024

**Authors:** Macarena G. GomezdelaTorre Clavel, Mason Youngblood, David Lahti

## Abstract

Domestication is the process by which species adapt to, and are artificially selected for, human-made environments. Few studies have explored how the process of domestication has affected the connection between behavioral traits and cognitive abilities in animals. This study investigated the relationship between personality and cognitive traits in domestic rabbits (*Oryctolagus cuniculus*). Fifteen individuals kept in a rabbit rescue facility were tested over a period of two months. We measured the linkage between behavioral traits (response to a novel object and exploration time) and cognitive performance. Our results suggest that there is no relationship between personality traits and problem solving abilities in domestic rabbits. In addition, our results suggest that exploration time is significantly repeatable at the individual level while latency to approach a novel object is not. Thus further research is needed to explore the relationship between cognitive and personality traits in domestic rabbits.

## Introduction

Domestication is a gradual and complex developmental and evolutionary process that has resulted in important morphological, physiological, and behavioral changes in animal species adapted to live in human-made environments (Price, 1984, 1999; Kaiser et al., 2015). Among the typical behavioral changes observed in domesticated species are differences in the frequency and magnitude of certain behaviors. For instance, domestication commonly leads to a decrease in aggression and exploratory behavior, and an increase tolerance to humans and conspecifics (Price 1984, 1999; Kaiser et al., 2015, Brust & Guenther 2015).

In recent years, many studies have explored how the personalities of domesticated animals differ from those of their wild counterparts (Range, Möslinger, & Virányi, 2012; Benhaim et al., 2013; Kaiser et al., 2015; Marino, 2015; Griffin, Guillette, & Healy, 2016; Brust & Guenther, 2017). Personality in non-human animals is defined as behavioral differences between individuals that are consistent over time and/or contexts (Carter et al., 2013; Biro & Stamps, 2008, Evans Ogden, 2012; Wolf & Weissing, 2012; Mackay & Haskell 2015). The behaviors that result in personality differences, such as boldness and aggression, are known as personality traits. During the 1990s, animal behavior researchers began to study personality traits such as boldness, exploration, predator avoidance, aggressiveness, and sociability (Carter et al., 2013). Correlated personality traits are called behavioral syndromes. For instance, the boldness-aggressiveness syndrome refers to the correlation of boldness and aggression in specific situations (Wolf & Weissing, 2012). Moreover, the continuum from boldness to shyness could be described as the differences in behavioral traits between individuals that are considered bolder, more aggressive, exploratory, and willing to take risks, versus those that are shy, unaggressive, less exploratory, and more cautious when taking risks (Sih & Del Giudice 2012; Oswald et al., 2012).

Cognition in animals can be defined as how animals process and use the information that they obtain from their environment to perform different functions, such as associative and social learning, memory, attention, self-recognition, and language development (Sih & Del Giudice, 2012; Shettleworth, 2000). Recently, there has been increased interest in the study of the link between cognition and personality traits in nonhuman animals (Griffin et al., 2015; Sih & Del Giudice, 2012). Some studies suggest that individuals have different ways to cope with environmental challenges and that this is related to a difference in cognition (Carere & Locurto, 2011; Koolhaas et al., 1999; Sih & Del Giudice, 2012). For instance, animals with proactive personalities are more aggressive, bold, neophilic, asocial, and active. Therefore, they are also more willing to explore and take risks. As a result, they might have more chances to interact with different environmental contexts and learn more quickly. In contrast, reactive individuals are non-aggressive, shy, neophobic, social, and inactive. In consequence, they are less willing to explore and take risks, and they might have less opportunity to explore novel environments and learn ways to cope with challenging situations and changing circumstances (Carere & Locurto, 2011; Koolhaas et al., 1999; Lermite et al., 2016). For instance, in guinea pigs bold and aggressive individuals learn faster than shy individuals (Guenther & Brust, 2017; Sih & Del Giudice, 2012). However, Guillete et al. (2015) found that slow-exploring black-capped chickadees (*Poecile atricapillus*) performed better in a learning task than fast-exploring conspecifics. This suggests that the direction of the relationship between personality and cognition is not homogeneous among species. (Doughtery & Guillette, 2018).

Domestic rabbits (*Oryctolagus cuniculus*) were domesticated only 1500 years ago (DeMello, 2010). A recent study demonstrated the existence of genetic and behavioral differences between domestic and wild rabbits that affect the development of their nervous system and behavioral repertoire (Carneiro, 2014). However, very few studies have focused specifically on the differences in behavioral traits among domestic rabbits, exploring only specific aspects of their personality such as boldness (Andersson et al., 2014). Other studies have used domestic rabbits to research the effect of hormones on behavior (Gosling 2008; Briganti et al., 2003). The relationship between differences in personality traits and variation in cognitive abilities in domestic rabbits has yet to be determined.

To explore the relationship between personality and cognition, we assessed consistency in personality traits along the bold-shy continuum and in cognitive abilities, such as problem solving and memory, in domestic rabbits. Consistency in total variation in personality traits within an individual is usually reported as repeatability (Falconer, 1981; Boake, 1989; Guenther & Brust, 2017). We then tested for a relationship between personality and cognition, following the methods of a study of guinea pigs by Brust & Guenther (2017). We made two predictions: that personality traits would exhibit considerable between-individual variation, and that bolder and more exploratory individuals would perform better in cognitive tasks than their shier and less exploratory conspecifics.

## Materials & Methods

### Experimental Animals & Housing

The study was carried out at the facilities of the Bunnies & Beyond Rabbit Rescue at Petsmart Flatiron in New York City. Following the protocol presented by Brust and Guenther (2017), a sample size of 15 individuals (9 females and 6 males) were tested to detect correlations between personality traits and cognition (Bell et al., 2009). The animals tested were adult rabbits of different breeds and ages that live individually or as bonded pairs. Individuals were housed singly or as pairs inside medium size steel stacked cages (38 in × 38 in × 37 in). Their enclosures contained one plastic tray and a bowl of fresh water available at all times. Each rabbit was fed ¼ cup of commercial rabbit pellets per day. Hay was given *ad libitum*. Standard operating procedures of the organization (i.e. food, health checks) were followed while conducting this study. These rabbits were surrendered, rescued from hoarding situations, or found as strays, and they remained under the care of Bunnies & Beyond until they were adopted.

### Ethical Standards

This study was conducted under the guidelines and approval of the Institutional Animal Care and Use Committee (IACUC) of Queens College of the City University of New York and under the United States Animal Protection Protocol#187.

### Personality - Open Field Test

To measure boldness in an unknown environment, each rabbit was placed in a novel arena (5 ft × 5 ft × 2 ft). The novel arena was divided into four quadrants, and a hideout (15 in × 15 in) was placed in the upper right corner of the arena. Each rabbit was placed in the lower left corner of quadrant one and allowed to freely explore the arena. The instantaneous sampling method was used to measure the movements and position of each rabbit every 15 seconds over 5 minutes (Altman, 1974; Andersson et al., 2014). Rabbits that spent a longer time exploring the novel arena were classified as bold while rabbits that remained a for longer time in the hideout were classified as shy (see Figure 1).

**Figure 1.**
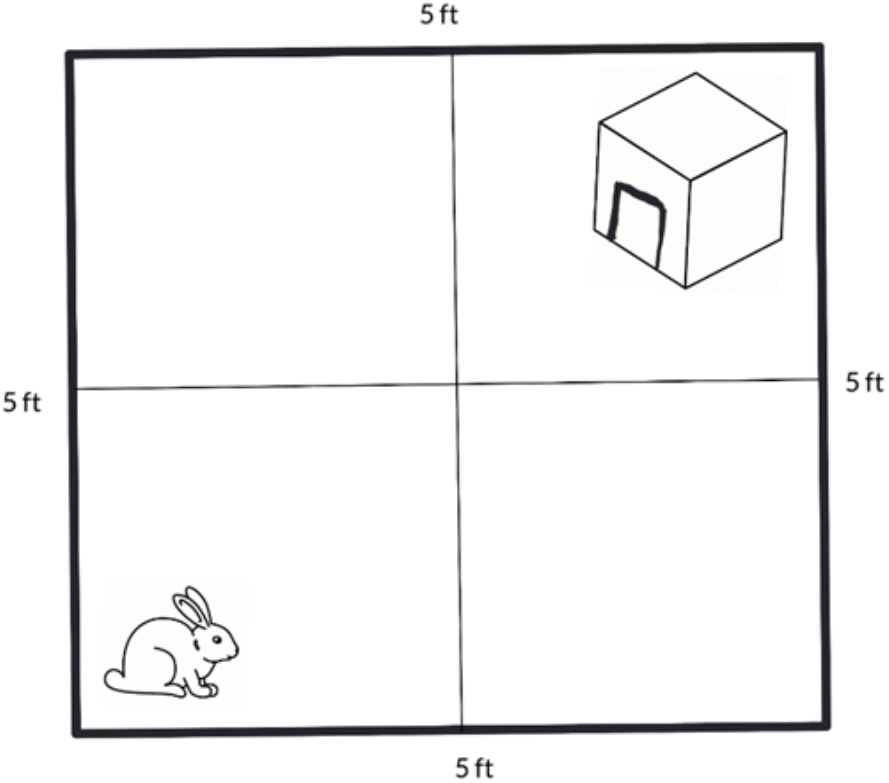
Quadrant divisions and hideout for the open field test.

### Personality - Novel Object Test

To measure boldness of each individual towards a novel object, each rabbit was placed in the novel arena before a novel object: a 10 cm high plastic toy. The object was placed in the center of the novel arena, and animals were observed for 5 minutes to measure the latency to approach and contact the object (see Figure 2).

**Figure 2.**
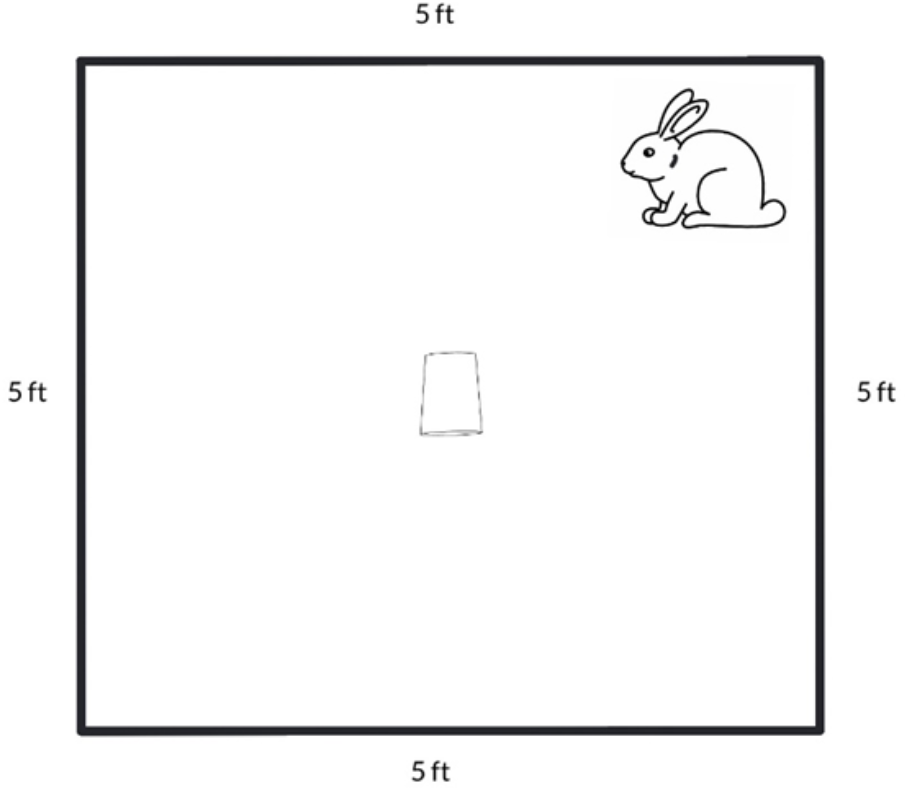
Novel object test with toy placed at the center of the arena.

Personality tests were conducted on the same day and repeated after one month with the same group of individuals to permit assessment of repeatability.

### Cognitive - Logic Board Test

A commercial logic board containing four compartments with lids was used for this test, and a piece of dried cranberry was hidden in one of the three compartments of the board. To solve the task, the rabbits had to open the lid of the compartment in which the treat was located to retrieve it. The test lasted 10 minutes. The latency to retrieve the hidden treat was recorded. Animals that were unable to solve the task were given the maximum latency score.

### Cognitive - Mage Labyrinth Test

A disposable cardboard maze labyrinth (34.5 in × 34.5 in × 10 in) was used for this test. The structure was divided into 8 compartments with panel holes (6.25 in tall × 5.5 in wide). This maze structure was built based on models that are commercially available at stores specialized in selling toys for rabbit pets, and it was replaced after every test to prevent odor contamination. The labyrinth was placed on the ground allowing sufficient space for unidirectional movement. The time to move through the labyrinth between the starting point and the ending point was recorded. The purpose of this test was to measure the rabbits’ ability to exit the labyrinth.

### Cognitive - T-maze Labyrinth Test

A disposable cardboard t-shaped maze labyrinth (34.5 in × 34.5 in × 10 in) was used for this test and replaced after completion. The purpose of this test was to measure cognitive functioning. The T-maze structure consisted of three segments: a start arm, and a right and left arm. The purpose of this test was to evaluate learning, spatial memory and spatial orientation. The rabbits were trained to run and enter the sidearm of the maze where a treat was located. The time necessary to reach the goal arm was recorded.

### Data Analysis

Eight of the 15 rabbits never retrieved the treat from the logic board, so that measure was recorded as a binary variable (0 = unretrieved; 1 = retrieved). Sex was also coded as a binary variable, with females as 0 and males as 1. Prior to analysis all binary variables were centered and all continuous variables were scaled and centered (Houslay & Wilson, 2017).

Repeatability of the two boldness measures was calculated using the *rpt*R package in R (Stoffel et al., 2017). *rpt*R uses MCMC generalized linear mixed modeling (GLMM) to calculate repeatability (R) using the following equation:

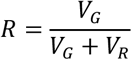

where V_G_ is the between-individual variance and V_R_ is the within-individual variance. High repeatability values indicate the consistency in the behavior within individuals and high variation among the behavior between individuals, which reveals animal personality traits (Boake, 1989; Brown & Shine, 2007; Schuster et al., 2017). Based on research in other small mammals, we classified rabbits that are prone to take risks and explore new environments as explorative and bold (Meijsser et al., 1989; Brust & Guether, 2015; Mazue et al., 2015). Parametric bootstrapping was used to estimate confidence intervals, and likelihood ratio and permutation tests to assess statistical significance (Stoffel et al., 2017). We ran separate univariate Gaussian models for the two boldness measures with 10,000 parametric bootstraps, 100,000 permutations, and individual identity as a random effect.

GLMM was conducted using the *MCMCglmm* package in R (Hadfield, 2010). Inverse gamma prior distributions with shape and scale parameters of 0.001 were used for all models, according to Guenther and Brust (2017). All models were run for 100,000 iterations with a burn-in period of 1,000 and a thinning interval of 10. Effects were considered to be statistically significant if the 95% highest posterior density intervals (HPDI) did not overlap zero. We assumed Gaussian distributions for both boldness measures as well as time to exit the labyrinth and time to retrieve food from the labyrinth, based on visual inspection of Q-Q plots with Kolmogorov-Smirnov confidence bands (see Appendix). We assumed a threshold distribution for retrieval from the logic board, as it was coded as a binary variable. Correlations between variables were calculated by dividing their covariance by the product of the square root of their variances, and then averaging across the MCMC chains to generate point estimates and 95% HPDIs (Houslay & Wilson, 2017).

To identify between-individual correlations between cognitive traits and boldness, a bivariate model was run with exploration time and latency to novel object as the outcome variables, sex, month tested, and the three cognitive measures as fixed effects, and individual identity as a random effect. In addition, in order to identify sex differences in cognitive traits, three univariate models were run with each cognitive measure as an outcome variable, sex as a fixed effect, and individual identity as a random effect. A single multivariate model with binary and continuous outcome variables could not be run due to convergence and mixing issues, but a separate binary model with only the two continuous variables (time to exit the labyrinth and time to retrieve food from the labyrinth) was run to assess whether they were correlated.

In all models the effective sample sizes for fixed effects were greater than 1,000, autocorrelations between successive means of fixed effects (accounting for thinning) were less than 0.1, and visual inspection indicated that the MCMC chains for the means and variances converged (see Appendix) (Hadfield, 2010).

## Results

The repeatability analysis indicates that exploration time was significantly repeatable at the individual level (*R* = 0.56; *p* = 0.012), whereas latency to approach was not (*R* = 0.36; *p* = 0.097). The bootstrap repeatability for both boldness measures at the individual level can be seen in Figure 3.

**Figure 3.**
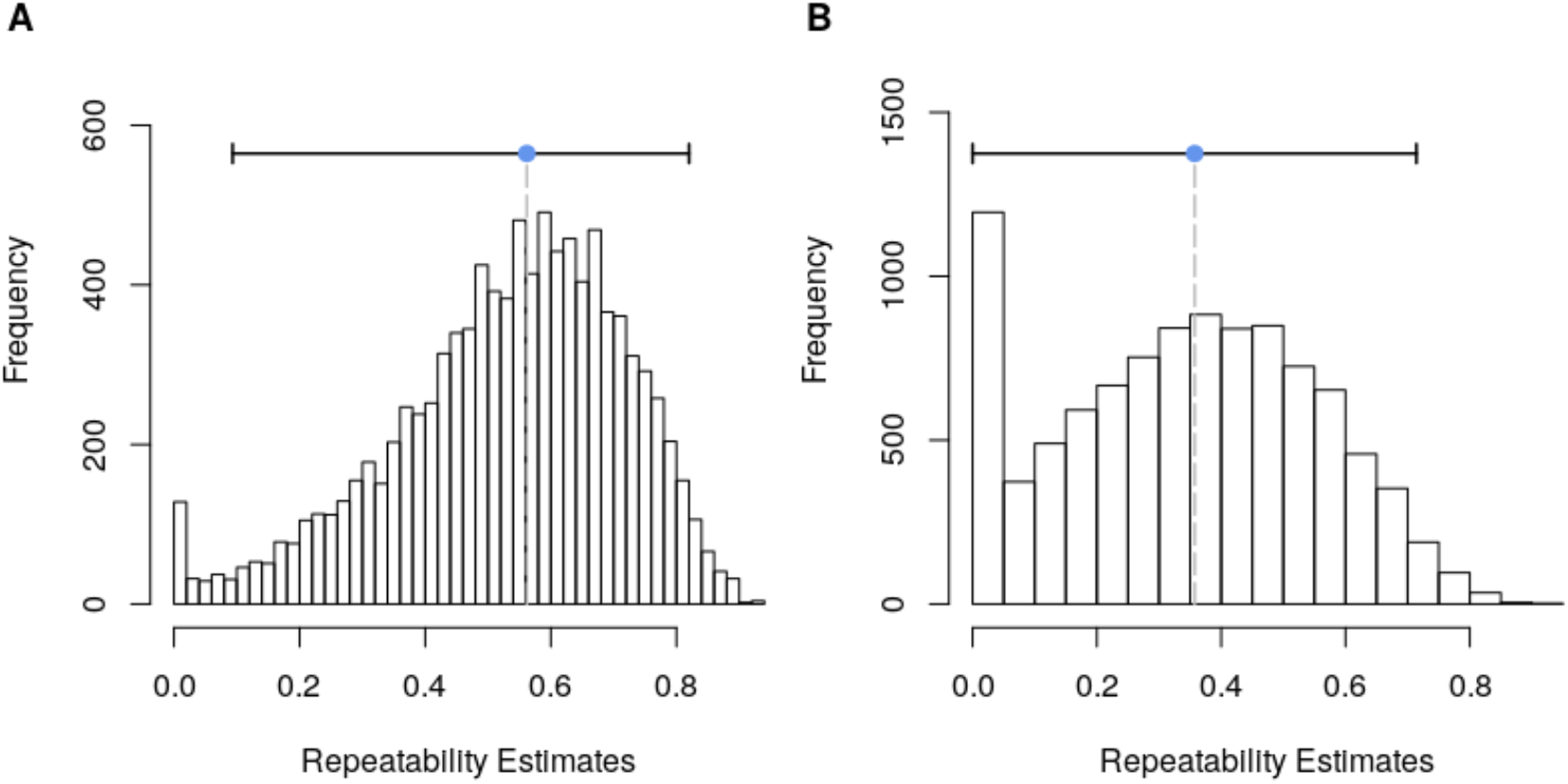
Bootstrap repeatability for (A) exploration time and (B) latency to novel object at the individual level. The blue line and dot indicate the point estimate for repeatability while the black bar indicates the 95% CI.

As seen in Table 1, month tested, sex, time to exit the maze labyrinth, and time to retrieve food from the T-maze labyrinth all had no significant effect on either exploration time or latency to novel object. Interestingly, retrieving food from the logic board appeared to negatively predict latency to novel object, but not exploration time (Table 1). Exploration time and latency to novel object were not significantly correlated with one another (*M* = - 0.54; 95% HPDI = [−0.95—0.0055]).

**Table 1.** The results of the bivariate model including side effects, the means of the posterior distributions, and the 95% HPDIs, and the *p*-values. Significant *p*-values are marked with asterisks (*<0.05).

|  |  | *Effect* | *M* | *95% HPDI* | *pMCMC* |
| --- | --- | --- | --- | --- | --- |
| Exploration Time |  | Month | -0.41 | [-0.90, 0.087] | 0.10 |
|  |  | Sex | 0.39 | [-1.076, 1.84] | 0.57 |
|  |  | Logic Board | 0.51 | [-0.74, 1.87] | 0.40 |
|  |  | Maze Labyrinth | 0.16 | [-0.53, 0.84] | 0.63 |
|  |  | T-maze Labyrinth | 0.13 | [-0.51, 0.76] | 0.68 |
| Latency |  | Month | 0.21 | [-0.39, 0.80] | 0.47 |
|  |  | Sex | -0.21 | [-1.52, 1.082] | 0.74 |
|  |  | Logic Board | -1.15 | [-2.34, -0.042] | 0.048 * |
|  |  | Maze Labyrinth | -0.014 | [-0.63, 0.59] | 0.96 |
|  |  | T-maze Labyrinth | -0.27 | [-0.83, 0.28] | 0.31 |

We found that there were no sex differences in cognitive traits (Table 2). Based on the results of the bivariate model, time to exit the labyrinth and time to retrieve food from the labyrinth were not significantly correlated with one another (*M* = 0.15; 95% HPDI = [−0.58—0.85]).

**Table 2.** The results of the three univariate models, including the fixed effects (outcome: fixed effect), the means of the posterior distributions, the lower and upper bounds of the 95% HPDIs, and the *p*-values.

| *Effect* | *M* | *95% HPDI* | *pMCMC* |
| --- | --- | --- | --- |
| Board Logic : Sex | 1.80 | [-1.47, 5.60] | 0.24 |
| Maze Labyrinth : Sex | 0.83 | [-0.22, 1.94] | 0.13 |
| T-maze Labyrinth Test : Sex | -0.43 | [-1.57, 0.73] | 0.44 |

## Discussion

The aim of the present study was to test the relationship between personality traits and cognitive performance in domestic rabbits. In contrast to other studies that have found a link between how bold and exploratory individuals are and their performance on problem-solving tasks (Carter et al., 2013; Brust & Guenther, 2017; Wat, Banks & McArthur, 2020), our results suggest that there is no relationship between personality traits and cognitive performance in domestic rabbits. Previous studies that reported a connection between boldness and cognitive performance have suggested that proactive individuals, which are more asocial, bolder and more willing to explore their environment, are more likely to learn faster because they have more chances to interact with novel environmental conditions than reactive individuals, which are more social, shier, and less exploratory (Bray et al., 2017; Brust & Guenther, 2017; Nawroth, Prentice & McElligot, 2016). However, a few studies show that the relationship between boldness and cognitive performance may vary according to the species and the context in which the problem-solving tasks occur (Schneider et al., 1991; Guillette et al., 2009, 2015; Albiach-Serrano, 2012; Trompf & Brown, 2013; Brust & Guenther, 2015; Bray et al., 2017; Lermite et al., 2016; Dougherty & Guillette, 2018).

In the case of domestic rabbits, there are several possible explanations for why our study failed to detect a relationship between personality traits and problem-solving performance. One possible explanation is that our results are explained by the effects of domestication on animal behavior (Brust & Guenther, 2015, Albiach-Serrano, 2012). Although implications of domestication in cognitive traits have not been extensively investigated, several studies have reported a weak link between personality and cognitive traits in domesticated species (Boissy, 2014; Brust & Guenther, 2015; Medina-García et al., 2017; Barnard et al., 2018; Dougherty & Guillette, 2018). Domesticated species have become adapted to artificial environments and low-risk conditions (Künzl et al., 2003; Kaisser et al., 2015). Domestication has removed selective pressures found only in natural habitats, such as predation and need for dispersion (Boice, 1973; Fox, 1967; Haase, 1980; Price, 1984; Ratner and Boice, 1975). As a result, the relationship between personality and cognitive performance in domesticated species may not be as evident as it is in the wild. For instance, studies done with guppies (*Poecilia reticulata*) show that bold individuals have a better cognitive performance than their shy conspecifics, which help them to identify predators and increase their chances of survival (Dugatkin and Alfieri, 2003). However, domestication seems to have contributed to the weakening or disappearance of the link between personality and cognitive traits in guinea pigs, budgerigars, and dogs (Brust & Guenther, 2015; Medina-García et al., 2017; Barnard et al., 2018). Artificial selection may have played a role in the dissociation of personality and cognitive traits in domestic lineages, as domestication has reduced the need for optimal performance (Brust & Guenther 2015). Artificial selection under domestication has also tended to favor less aggressive individuals that are better adapted to live in man-made environments and in close contact with humans (Brust & Guenther, 2015; Kaiser et al., 2015). Therefore, our data might indicate that the stability and safety of man-made environments caused a decrease in the sensitivity of domesticated rabbits to environmental changes, resulting in a weaker linkage between behavioral and cognitive traits.

Many studies have reported differences in behavior, morphology, and physiology between wild and domestic species. Some of these studies shown that domestic species, such as guinea pigs and rats, perform better in learning and memory tasks than their wild-counterparts (Kruska, 1998; Kaiser et al., 2015; Brusini I, Carneiro M, Wang C, et al., 2018). In a study that compared the anatomy of brain structures of domestic and wild rabbits, a reduction of the size of the amygdala was observed in domestic rabbits. The amygdala is a brain structure involved in the processing of emotional memory and the triggering of the fight-or-flight response (Brusini I, Carneiro M, Wang C, et al., 2018). Similarly, a higher level of activity was detected in the medial prefrontal cortex (mPFC) of domestic rabbits. The mPFC supports the processing of social information, and along with the hippocampus, it is involved in rapid learning and memory consolidation in humans, monkeys, and rodents (Grossmann, 2013; Euston, Gruber, & McNaughton, 2012). Studies on rabbits show that the mPFC facilitates associative learning (Buchanan et al., 1994). However, cognitive performance may depend not only on brain morphology but also on a variety of factors such as environmental conditions and the context of cognitive tasks. For instance, domesticated and wild gerbils born in captivity performed better in an auditory discrimination learning task than wild gerbils living in their natural habits (Kaiser et al., 2015). This makes it difficult to predict a particular difference in cognitive performance based only on personality traits (Kaiser et al., 2015; Brust & Guenther, 2015; Dougherty & Guillette, 2018). Consequently, domestication could explain the lack of the association between personality traits and cognitive performance in domestic rabbits.

Another possible explanation for the lack of relationship between boldness and fast problem-solving performance is that domestic rabbits that live in enriched human-made habitats have lower levels of stress (Trocino & Xiccato, 2010; Trocino et al., 2013), which may help them to perform better in cognitive tests. This idea is supported by studies done in dogs, pigs, rats and rhesus monkeys. Dogs, pigs, rats, and primates reared in enriched conditions are better problem solvers than individuals reared in deprived environments (Sacket, 1972; Schneider et al., 1991; Asher et al., 2016; Barnard at al., 2018). The 15 rabbits we tested lived in large size cages in which they had toys, and water and food *ad libitum*. In addition, they were scheduled to spend at least two hours per day playing in a puppy play pen with tents and tunnels and under the constant stimulation and care of the volunteers of the rescue, which may help them to socialize and bond with conspecifics and humans. This might explain the observed lack of relationship between boldness and better performance in cognitive tasks. Further studies on the relationship of personality and cognition in animals could help us to explore and understand how domestication may have altered the linkage between personality traits and cognitive performance in domestic rabbits, and how this species adapted to novel artificial environments.

Our second finding was that exploration time is significantly repeatable at the individual level. This suggests consistency of personality traits over time (Brust & Guenther 2015; Koski, 2014). A value of 56% of variation in individuals (*R* = 0.56) is within the range of the majority of repeatability estimates for behaviors in species of diverse taxa. High repeatability in exploration time suggests consistency in the behavior within individuals, which may be considered a personality trait in domestic rabbits (Bell et al. 2009). A study in fish and avian species found that exploratory behavior and boldness are important for dispersal (Mazue et al., 2015; van Oers et al., 2004). If this is so, then understanding variation and consistency in boldness within populations may be important for conservation of re-introduced or translocated species (Bremner-Harrison, 2004). In the case of rabbits this is relevant because they have become an endangered species in Europe (Price, 1984; Virgos et al., 2006). Domestic rabbits were not consistent in approaching a novel object, which suggests that this personality trait is not stable over time. Other studies in rabbits have found that higher repeatability in the novel object test. However, values varied when the novel object test was performed in familiar versus novel environments (Andersson, 2014). The repeatability observed in the present study might also be explained by the absence of control for features such as individual breed, age, past life experience, and environmental conditions. The difference in repeatability in our results suggest that exploration time and novel object behaviors reflect different personality dimensions in rabbits (Stamps & Groothuis, 2010; Andersson, 2014). Further research is required to investigate the relationship between personality traits involved in exploration and approach to novel objects, and their impact on cognitive performance in domestic rabbits.

We did not observe any sex difference in cognitive traits. Some other studies have reported a significant difference in levels of anxiety and boldness between young rabbits of different breeds and sexes. However, sex differences in personality traits are no longer significant by the time rabbits become adults (Andersson, 2014). Future studies should focus on investigating how breed, sex, and age affect personality traits and their association with cognitive performance in domestic rabbits.

To our knowledge the current study is the first to examine the relationship between personality and cognitive traits in domestic rabbits. Even though our results indicate a lack of correlation of personality traits and cognitive performance, the small sample size in this study highlights the need for further research with a larger number of individuals. Small sample size has often hindered researchers’ ability to investigate reasons for variation in personality and its relationship with performance in problem-solving tasks (Carere & Locurto, 2011; Andersson, 2014). Additionally, we propose controlling for other features, such as the individual breed, age, past experience, housing conditions, and enriched environments. These features could potentially affect both personality and cognitive traits. Therefore, future researchers should consider the influence of these variables on the relationship between personality and cognitive performance. Also, in order to better understand the effect of domestication on personality and cognitive traits, I emphasize the need of designing new tasks to test both personality and cognition in domestic rabbits.

In conclusion, I found no relationship between personality traits and cognitive performance. This is consistent with previous findings of lack of association between behavioral and cognitive traits in species that have undergone domestication and artificial selection. Exploration time of a novel arena, which is related to boldness and dispersal, was found to be repeatable in domesticated rabbits, which may indicate a personality trait in the species. However, latency to approach a novel object was not repeatable. No sex differences were found in cognitive traits. These results demonstrate the importance of learning more about how domestication and human contact may have influenced the relationship between personality and cognitive abilities in domestic rabbits.

## Data Availability

The data and R code are available in the Harvard Dataverse repository: https://doi.org/10.7910/DVN/WOZ8AQ

## Acknowledgments

We would like to thank Cindy Stuts and Erin Alana from Bunnies & Beyond, Dr. Levison VMD from Symphony Vet Center, and all members of the Lahti lab.

## Appendix

### Distribution Identification

**Figure S1.**
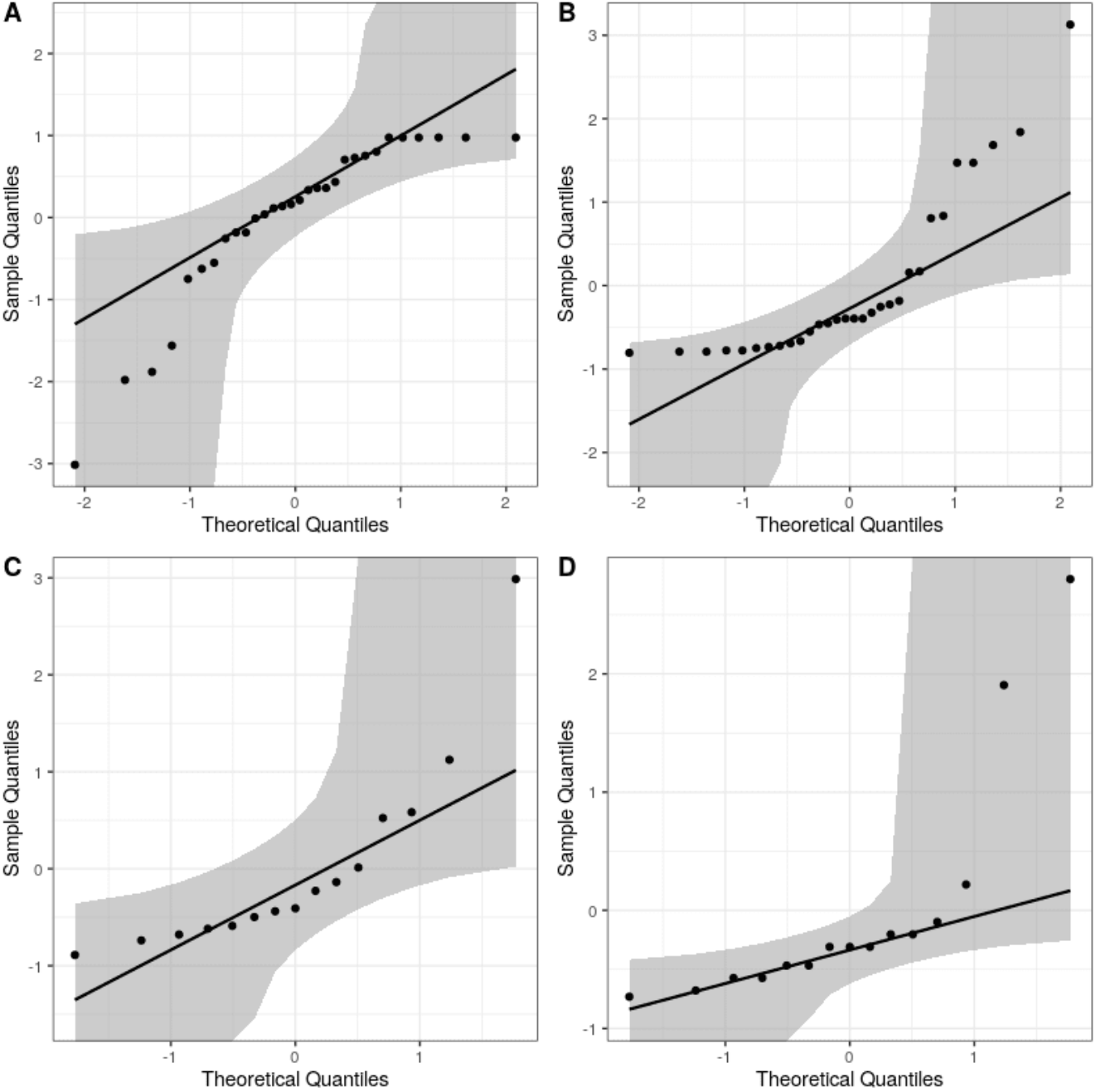
Q-Q plots with the expected values according to a gaussian distribution in black, and the Kolmogorov-Smirnov confidence bands in grey, for: (A) exploration time, (B) latency to approach a novel object, (C) time to exit the labyrinth, and (D) time to retrieve food from the labyrinth.

Q-Q plots were constructed using the *qqplotr* package in R (Almeida et al., 2017). Kolmogorov-Smirnov confidence bands were used because they correspond to a statistical test, but we should note that confidence bands constructed with more conservative techniques (e.g. bootstrapping and tail-sensitivity) do not contain all points.

### MCMC Diagnostics

**Figure S2.**
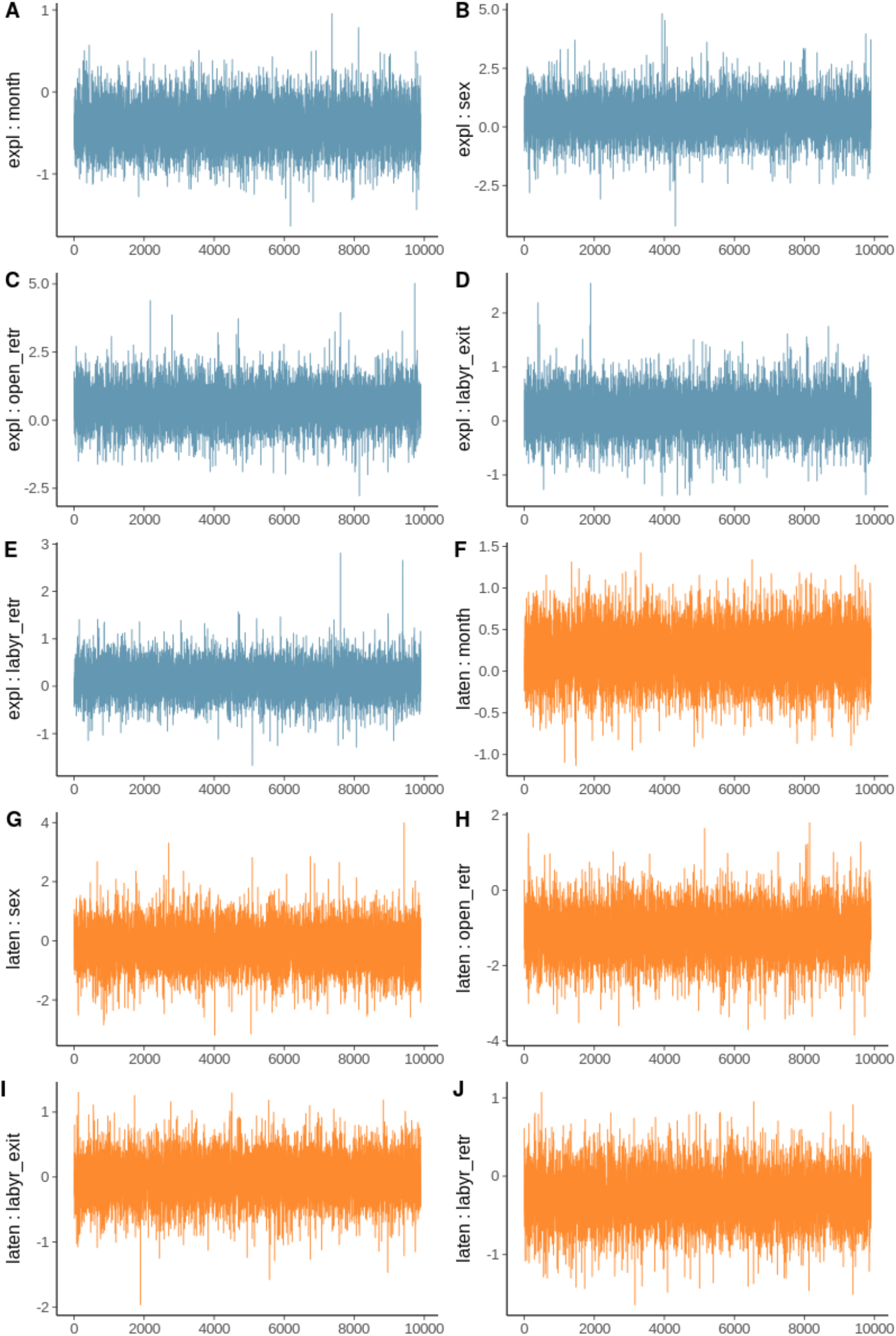
MCMC traces for the bivariate model. Blue traces correspond to the effects of the fixed effects on exploration time, while orange traces correspond to the effects of the fixed effects on latency to novel object.

**Figure S3.**
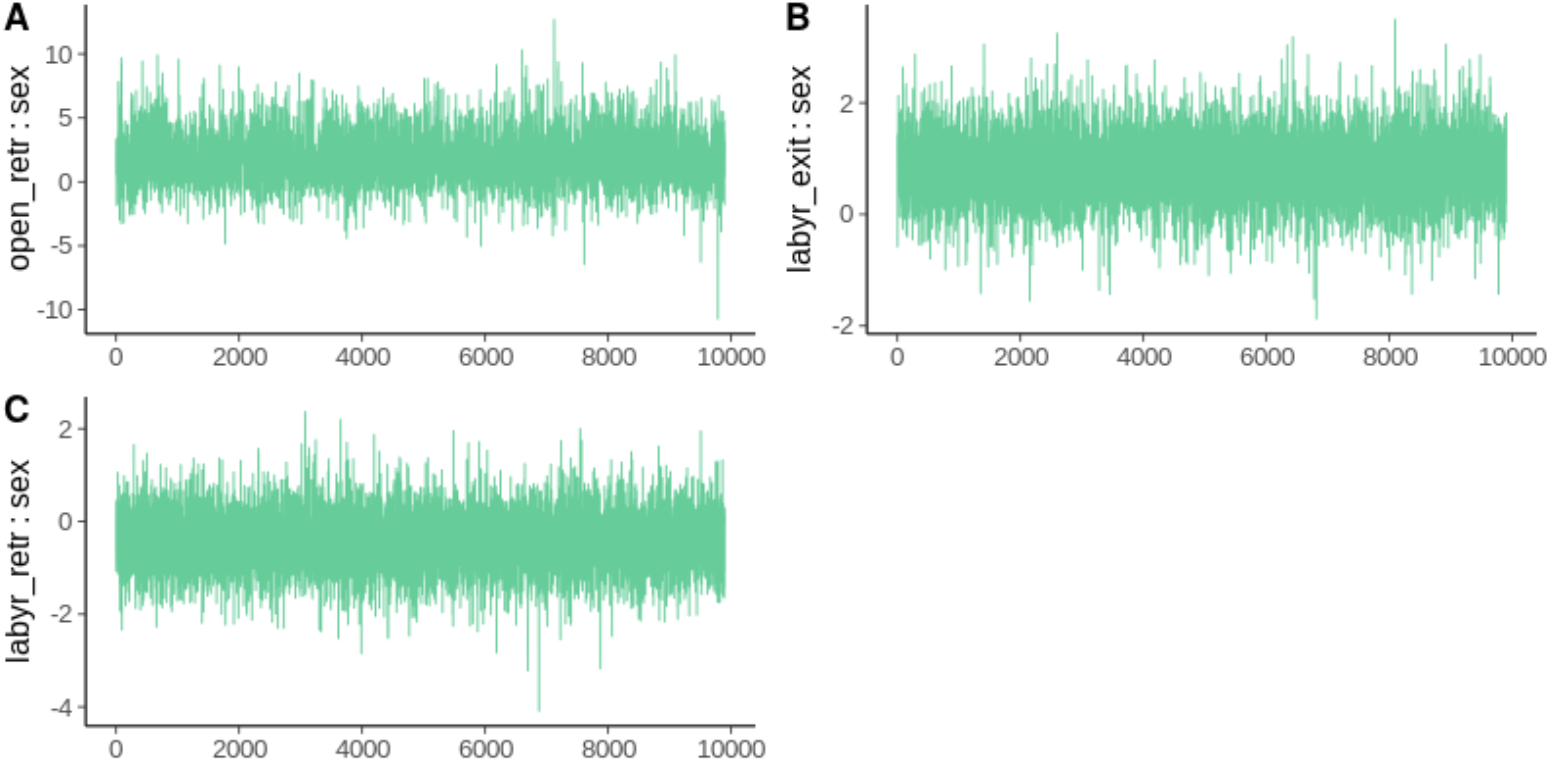
MCMC traces for the univariate model.

